# Cell fate-decision as high-dimensional critical state transition

**DOI:** 10.1101/041541

**Authors:** Mitra Mojtahedi, Alexander Skupin, Joseph Zhou, Ivan G. Castaño, Rebecca Y. Y. Leong-Quong, Hannah Chang, Alessandro Giuliani, Sui Huang

**Affiliations:** Department of Biological Sciences, University of Calgary, 2500 University Dr. NW, Calgary, AB T2N 1N4, Canada; Institute for Systems Biology, 401 Terry Ave N, Seattle, WA 98109, USA; Luxembourg Centre for Systems Biomedicine, 7, Avenue des Hauts-Fourneaux L-4362 Esch-sur Alzette, Luxembourg; 5AM Ventures, 2200 Sand Hill Road, Suit 110, Menlo Park, CA 94025; Corporación Parque Explora, departament of innovation and design, Colombia; Istituto Superiore di Sanità, Roma, Italy

## Abstract

Cell fate choice and commitment of multipotent progenitor cells to a differentiated lineage requires broad changes of their gene expression profile. However, how progenitor cells overcome the stability of their robust gene expression configuration (attractor) and exit their state remains elusive. Here we show that commitment of blood progenitor cells to the erythroid or the myeloid lineage is preceded by the destabilization of their high-dimensional attractor state and that cells undergo a critical state transition. Single-cell resolution analysis of gene expression in populations of differentiating cells affords a new quantitative index for predicting critical transitions in a high-dimensional state space: decrease of correlation between cells with concomitant increase of correlation between genes as cells approach a tipping point. The detection of “rebellious cells” which enter the fate opposite to the one intended corroborates the model of preceding destabilization of the progenitor state. Thus, “early-warning signals” associated with critical transitions can be detected in statistical ensembles of high-dimensional systems, offering a formal tool for analyzing single-cell’s molecular profiles that goes beyond computational pattern recognition but is based on dynamical systems theory and can predict impending major shifts in cell populations in development and disease.

## Introduction

A multipotent stem cell or progenitor cell is in a state that poises it to commit to one of two or more predestined lineages and to differentiate. Yet, its state-characteristic gene expression profile in the high-dimensional gene expression state space is robustly maintained because the cell is in a stable attractor state [1,2] (“ground state” [3]) of the gene regulatory network (GRN). Thus, commitment to a lineage involves overcoming this stabilization as genes alter their expression in a coordinated manner to establish the new gene expression pattern that implements the new phenotype of the differentiated state (Fig 1). Individual cells can, due to noisy gene expression fluctuations, transiently approach the border of the attractor in one or several dimensions and thereby be transiently “primed” to exit the basin of attraction, and by chance or biased by external conditions, differentiate into one of the predestined lineage accessible to the respective multipotent progenitor [4,5,6,7]. However the fundamental question remains whether differentiating cells exit the progenitor attractor simply by harnessing rare chance configurations of expression of the appropriate regulatory proteins to “jump” into a new nearby stable attractor state [8,9,10,11] or instead, by undergoing a larger-scale destabilization of their (high-dimensional) gene expression state [12,13,14].

**Fig 1.**
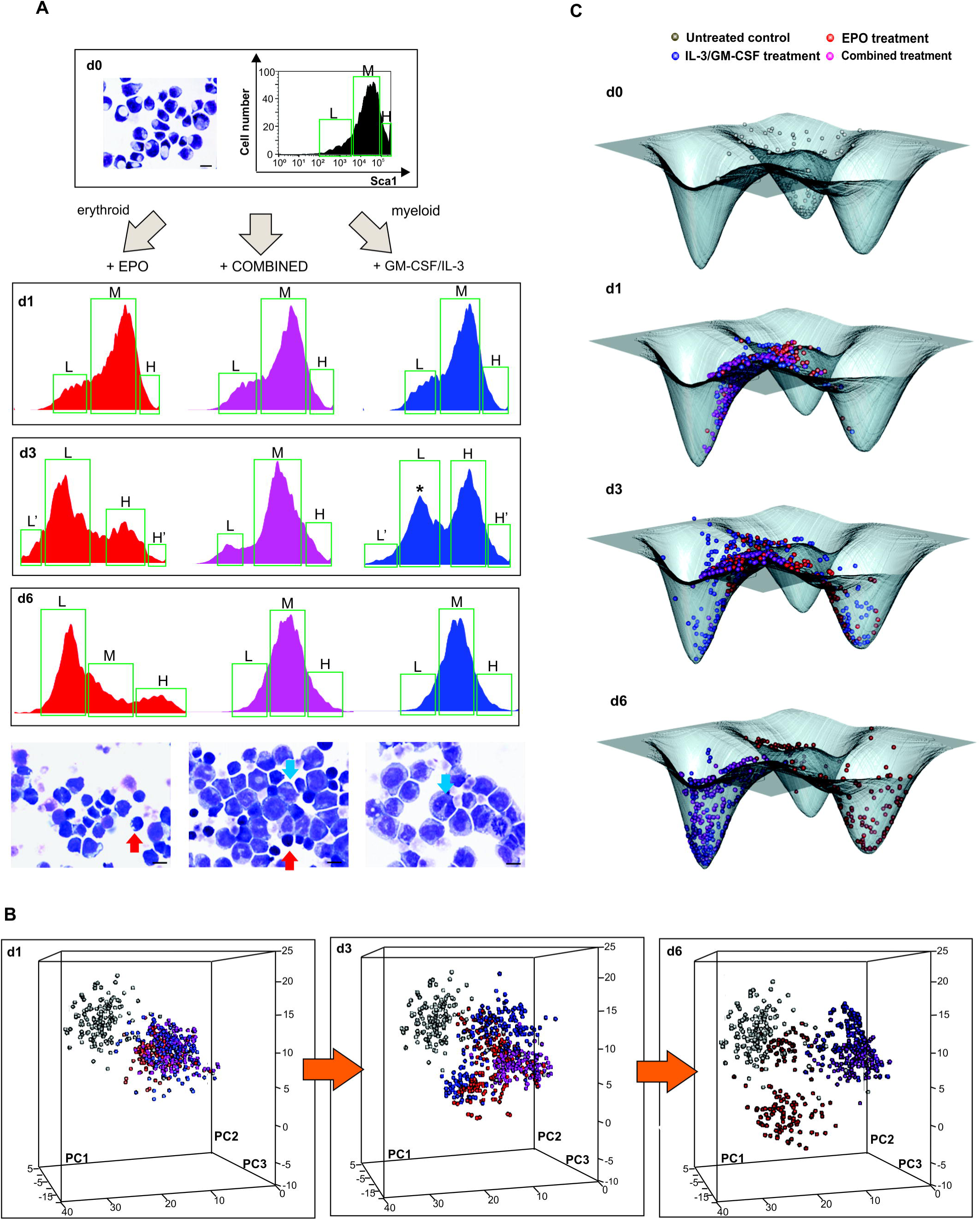
Single-cell analysis of transcript expression during binary fate decision in EML cells. **(A)** A progenitor EML cell population was stimulated with EPO (left), IL-3/GM-CSF (right) or with a combination of EPO, GM-CSF/IL-3 (center). Flow cytometry histograms of Sca1 surface expression were gated into Sca1^LOW^ (L), Sca1^MEDIUM^ (M) and Sca1^HIGH^ (H) fractions or subpopulations (green boxes) during FACS sorting of single cells at the indicated days for use in later analysis (Fig 2). At d3, further division to account for the extreme outliers (L’, H’)* indicates “rebellious cells” (see text). **(B)** For visualization of individual cells’ transcript expression patterns (of the *m*=17 genes) cells were projected onto a dimension-reduced state space spanned by the three first principal components (PC) following principal component analysis (PCA, see S1 Appendix). Each sphere represents a cell, colored according to treatment: untreated progenitors (grey); cells treated with EPO (red), cells treated with GM-CSF/IL-3 (blue); and combined-treated cells (purple). **(C)** To calculate a quasi-potential landscape for the three cell types, a Gaussian filter with s =2 was applied to PC1 and PC2 coordinates of cells at d0 and d6 treated with EPO and GM-CSF/IL-3 leading to a smooth 2-dimensional distribution p. With the (quasi-)steady state assumption [15], the attractor landscape was visualized relative to a base level of 0 by −log(p +1).

A cell fate choice and fate commitment driven by a destabilization of the progenitor attractor state until cells “spill out of it” would represent a critical state transition [15,16]. Herein, a stable attractor state of a system is gradually destabilized due to a steady and monotonical change in one characteristic of its underlying control structure (a systems parameter) until the system suddenly transits a “tipping point” (bifurcation) at which the stable attractor state disappears and other attractors become accessible. The progenitor cells in that destabilized attractor would then move to the(se) new stable state(s) that represent the gene expression pattern of new cell phenotypes. This formal description would explain multi-potentiality and the quasi-irreversible lineage restriction beyond a “point of no-return” [14].

Critical transitions of a system (like a cell) can occur because of the presence of non-linear interactions between its underlying component parts (genes, proteins) that collectively produce multiple distinct potential behaviors (cell phenotypic states) and if the realizable range of the value of critical parameters that characterize these interactions encompass qualitatively distinct behavioral regimes. Even if specific details of the interactions and the identity of the critical parameter are not known, stochastic fluctuations or certain perturbations can expose a system’s approach towards a critical state transition. This is manifest as “early warning signals” and is essentially a consequence of the destabilization of an attractor state that precedes the bifurcation event. Early warning signals can be exploited to predict a qualitative system-wide shift in a complex nonlinear system, as has been applied to material properties, ecosystems, social systems and disease states [15,16,17,18].

The principle of a bifurcation governing cell fate choice naturally unites the two classical models of binary cell fate decision of multipotent progenitor cells at developmental branch-point: the stochastic (intrinsic) and the deterministic (instructive) models [19,20,21,22,23]. In the stochastic model [24,25] the cells randomly assume a (pre)committed, or primed state that renders them responsive to the fate-specifying growth factor which then acts to selectively expand (“select”) these primed cells which in turn would proliferate and terminally differentiate. In the deterministic model [26,27], the same factors act to specifically instruct the cell which gene to turn on and off to establish the gene expression pattern of the prospective fate. These two models are not mutually exclusive and experimental evidence support either scheme depending on experimental design [19,20,21,22,23].

Here we use single-cell gene expression analysis to examine in a model system the fate commitment of blood progenitor cells either into the erythroid cells (precursors of erythrocytes), promoted by the growth factor (cytokine) erythropoietin (EPO), or into the myeloid lineage (precursors of monocytes and granulocytes), promoted by the cytokines GM-CSF and IL-3. We show that the formalism of critical state transitions, so far only demonstrated for examples in which the system state is characterized by one variable [17], can (*i*) be applied to a high-dimensional system, namely the gene expression pattern defining a mammalian cell state, while (*ii*) at the same time, taking advantage of the fact that the system is present in an ensemble of replicates: a population of cells. To this end, we introduce a new index *I*_*C*_ computed from high-dimensional single-cell gene expression profiles of cell populations and show that it can serve as an early warning signal for an impending cell state transition. Critical transitions also explain the long-observed phenomenon of “rebellious cells” that differentiate into the direction opposite to that instructed by the growth factors. Single-cell resolution analysis of cells exposed to conflicting stimuli also confirm that developmental trajectories are robust and predestined, as predicted by early models [14,28], and that the stochastic and deterministic scheme of cell fate control coexist.

## Results and Discussion

### 1. Single-cell gene expression patterns during binary cell fate decision

To determine if differentiation goes through a tipping point in high-dimensional gene expression state space we studied the commitment of the murine multipotent hematopoietic precursor cell line EML into an erythroid and myeloid fate when stimulated by EPO and GM-CSF/IL-3, respectively [4]. We also treated cells with a combination of EPO and GM-CSF/IL-3 to separate a generic destabilization from the specific fate choice because we reasoned that the latter should be neutralized by the combination treatment. To ensure that heterogeneity of the starting cell population is due to dynamic fluctuations and not to pre-existing pre-committed cells (which would merely be selectively enriched by the respective growth factors) we used a cell line, as opposed to primary cells, that allows for the study of populations recently derived from a single common ancestor. We monitored transcript expression patterns at single-cell resolution using qPCR to harness the information provided by a statistical ensemble of (randomly distinct) cells which manifests the stability of a given nominal cell state.

Exit from the progenitor state was first monitored by flow cytometry measurement of the down-regulation of the stem-cell markers Sca1 and c-kit. The induction of a bimodal distribution with a new discrete subpopulation with lower Sca1 (as well as c-kit) surface protein expression confirmed the switch-like state transition (Fig 1A). Fig 1B shows the time course of single-cell transcript patterns of 19 selected genes (listed and explained in S1 Fig and S1 Table) known to be functionally involved in or to mark fate commitment of the EML cells, visualized by plotting each cell into the Cartesian space spanned by the three principal components (PC) following principal component analysis (PCA) to reduce the 19-dimenional state space (Appendix A Supplementary Methods). The “cloud” of untreated cells (black, depicted for reference for each time point) spreads upon treatment (colored balls), reaching highest diversity at day 3 (d3). The cells then coalesced into two distinct dense clusters at d6 (blue and red) representing the erythroid (red) and myeloid (blue) lineages which were identified by the characteristic expression of erythroid or myeloid transcript levels (S2 Fig and S1 Table). As shown in S3 Fig, in this single-cell qPCR, measurement noise was only a small fraction of the biological cell-to-cell variability, thus the dispersion of points in state space largely reflects the biological diversity of cells. Loading of gene scores show that PC1 captures the erythroid-myeloid dichotomy, whereas PC2 reflects the stemness-differentiation axis (S4 Fig). Single-cell measurement also provides the local cell density for each position in state space which can be visualized as the elevation of an approximate quasi-potential landscape [12] (Fig 1C, legend) which shows the three attractor states as minima.

Intriguingly, at d3 some cells consistently went in the “wrong” direction, opposite to the instruction by the cytokines (e.g. some EPO-treated cells were associated with the myeloid cell cluster and vice versa). These “rebellious cells” disappeared at d6 – either by “transdifferentiating” to the “correct” lineage or by dying out (see below). Progenitor cells receiving a combined treatment also diverged at d3 but stayed in an intermediate “undecided” region of the state space before joining the myeloid cluster (Fig 1B). Thus, the conflicting signals delayed the fate decision but ultimately a uniform decision is made. This behavior corroborates the notion that gene expression change during lineage determination is not simply instructed by external growth factors, but also governed by intrinsic constraints that channel cells towards predestined attractors of the GRN and do not allow for stable intermediates, as Waddington first predicted [28]. In this case it appears that the attractor for the myeloid fate is more readily accessible.

### 2. High-dimensional critical state transition in ensemble of systems

Independent of the (unknown) detailed dynamics of the underlying GRN, a destabilization and disappearance of even a high-dimensional attractor state is a bifurcation event and should display the signatures of an approach to a critical phase transition [16] at which cells would undergo a discontinuous switch towards the destination state. While the bimodal distributions of Sca1 (Fig 1A) after d3 already suggest a quasi-discrete transition, they cannot reveal a destabilization in a high-dimensional state prior to the switch. Recently reported cases of critical transitions in stressed ecosystems and disease processes [refs. in [17]] pertain to low-dimensional systems in which typically one systems variable was observed longitudinally. By contrast, here we examine time snapshots of states of a high-dimensional system (19-dimenional cell state vector) embodied by the GRN.

From theoretical consideration, a critical destabilization and transition to a new attractor will be manifest in two changes in the correlation statistics (as explained and derived in S2 Appendix): First, a decrease of cell-cell correlation *R*(cell *k*, cell *l*) between all pairs of the *n* cell state vectors in the *m*=17-dimensional gene space; this reflects the expected increase of amplitudes of random fluctuation of gene expression due to the weakening “attracting force” in the “flattening” basin of attraction prior to the bifurcation [29,30]. Second, a concomitant increase of gene-gene correlation *R*(gene *i*, gene *j*) between all pairs of “gene vectors” that describe the gene expression values of each gene across all the cells; this corresponds to the increase of long-range correlations of state variables in time and/or space described in many phenomenological analyses of critical state transitions [17]( *9*)( *9*). The overall increase in the correlation between the gene vectors arises because of the symmetry-breaking destabilization and is plausible from two different perspectives: (i) as a consequence of the “range restriction effect” of correlation in statistics when the dominance of the symmetric stochastic fluctuations in the attractor yields to non-symmetric, regulated change of gene expression [31,32] or (ii) as a consequence of the appearance of a saddle-node in the dynamical system description through which the individual cells pass. A detailed mathematical derivation of is provided in the S2 Appendix. This reasoning motivates an index for critical transitions, *I*_*C*_:

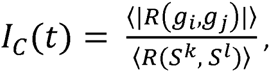

where *g* are gene vectors and *S* are the cell state vectors at sample time *t* and 〈*R*(…,…)〉 denotes the average of all Pearson’s coefficients of correlation. We postulate that *I*_*C*_ increases towards a maximum when cells go through the critical state transition (see S2 Appendix). Recently, Chen et al. proposed a similar index for full transcriptome time courses which for lack of single-cell resolution state vectors estimates state diversification differently and involves the prior computational selection of a subset of genes in the same data [30].

Fig 2A shows the *n*×*n* heat map for cell-cell correlation coefficients *R*(*S*^*k*^, *S*^*l*^) for all pairs of the *n*=1600 cells for the three treatments (EPO, GM-CSF/IL-3 and combined) at each time point *t*. The diagonal shows that correlation of cells within the populations decreases at d1 and notably at d3, compared to d0, and increases again at d6, indicative of a transient diversification of cell states and a return to a more homogenous population consistent with an attractor state. Since we also recorded the cells’ position with respect to the Sca1 surface marker expression (roughly partitioning the population into three fractions, Sca1-high (*H*), Sca1-medium (*M*) and Sca1-low (*L*) – see Fig 1A) one can see that the decrease of correlation was not due to comparing cells across subpopulations in bimodal populations (Fig 1A). The higher correlation among the cells within the extreme-low Sca1 fraction (*L’*) in both EPO and GM-CSF/IL-3 treatment is consistent with advanced commitment of cells which are enriched in the Sca1-low fraction towards the erythroid fate as previously reported [4]. By contrast, the high correlation among the *H* cells at the end of EPO treatment reflects the “rebellious” cells that became myeloid under EPO treatment.

**Fig 2.**
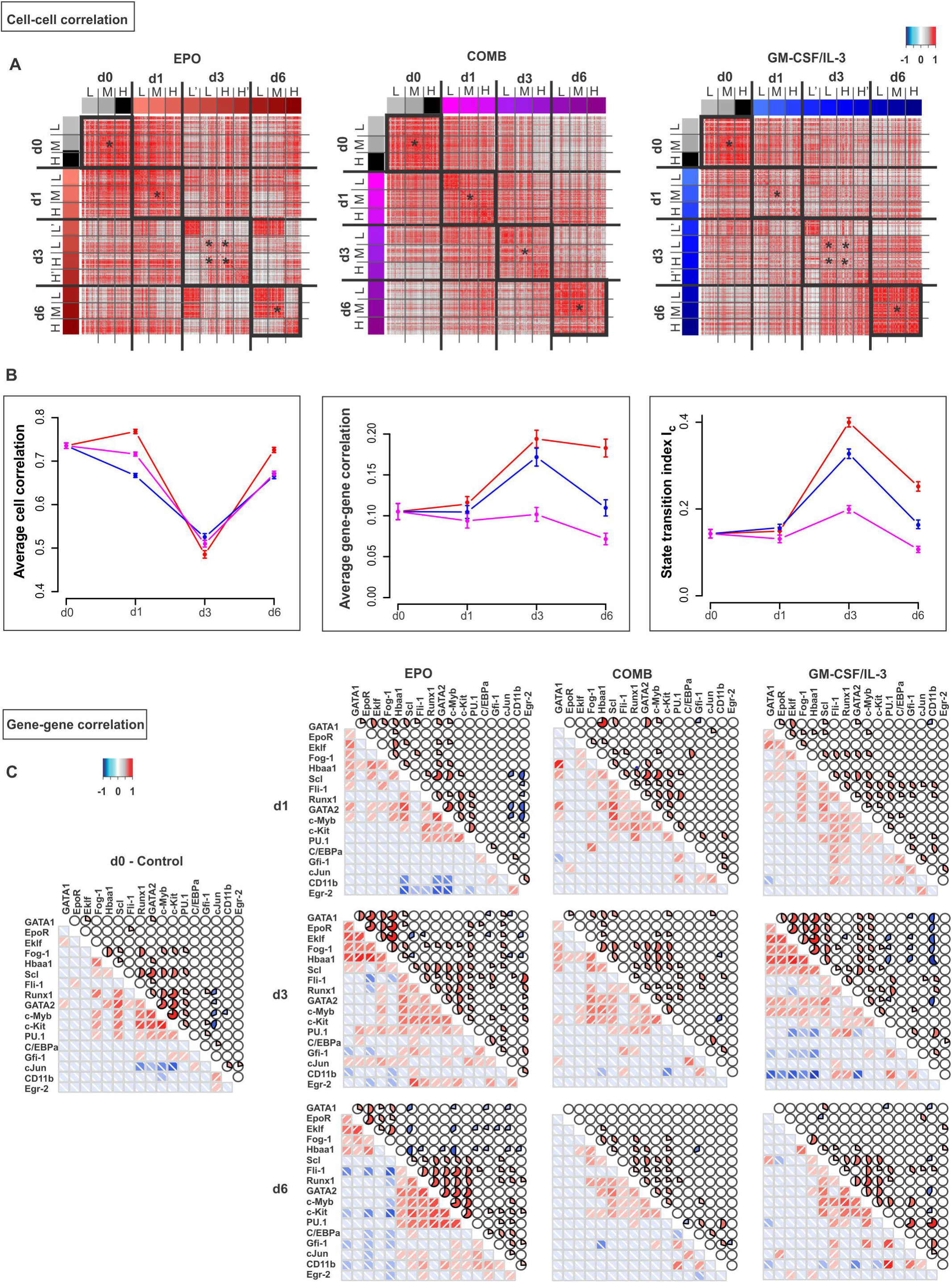
Critical transition during lineage commitment. **(A)** Cell-cell correlation matrices displaying the Pearson correlation coefficient *R*(*S*^*k*^, *S*^*l*^) for all pairs of cells in states *S*^*k*^ and *S*^*j*^ (see S2 Appendix). *R* calculated for a set of 150 progenitor cells, 500 EPO-treated, 500 GM-CSF/IL-3-treated and 450 combination-treated (COMB) cells from data used in Fig 1. Black squares (diagonal) emphasize the higher correlation between cells within the nominally same population. Two control genes (GAPDH and TBP) were excluded from this analysis. L’, L, M, H, H’ indicate the Sca1 fractions shown in Fig 1: extremely low, low, medium, high and extremely high level of Sca1 expression, respectively. **(B)** Average Pearson correlation coefficients for all cell-cell pairs (left) and all gene-gene pairs (center) as well as the state transition index *I*_*c*_ = 〈|*R*(*g*_*i*_, *g*_*j*_)|〉/〈*R*(*S*^*k*^, *S*^*l*^)〉 at various time points. Cell-cell correlation coefficients were calculated for the central fractions/subpopulations in panel A(*). Error bars indicate SEM. **(C)** Gene-gene correlation matrices for the 17 genes of interest and the two endogenous control genes for the three treatments at various time points where correlation is indicated either by color (lower matrix triangle) or solid color segment in pie chart. Color values for magnitude of correlation coefficient for both matrices (A, C) are shown in color bar.

The second criterion for a critical state transition, the increase in gene-gene correlation 〈*R*(*g*_*i*_, *g*_*j*_)〉, between the genes is shown in Fig 2B. Both EPO and GM-CSF/IL-3 treatment resulted in almost a doubling of 〈*R*(*g*_*i*_, *g*_*j*_)〉 at d3 which returned towards baseline at d6. The heat-maps of the raw data (Fig 2C) show that the increase of 〈*R*(*g*_*i*_, *g*_*j*_)〉 resulted from correlated (red) as well as negatively correlated gene pairs (blue) at d1, but more pronounced at d3. By contrast, genes were mostly uncorrelated in the progenitor state, consistent with the dominance of random fluctuations around the attractor state.

Together, the cell-cell and gene-gene correlation gave rise to a temporal course of the index *I*_*C*_ that sharply peaked at d3 after induction of either fate commitment, which thus marks the critical transition and coincides with lineage separation in state space (Fig 1B).

### 3. Alternative monitoring of myeloid commitment reveals rebellious cells

To exclude that the gene-gene and cell-cell correlation behavior is an idiosyncrasy linked to monitoring the exit from the progenitor attractor along the direction of Sca1 reduction, we also monitored and dissected differentiation along the axis of increase of the differentiation marker CD11b, a reliable indicator of myeloid differentiation (Fig 3A). Following GM-CSF/IL-3 treatment, CD11b surface expression first increased and then Sca1 decreased, from CD11b^LOW^/Sca1^HIGH^ to CD11b^HIGH^/Sca1^LOW^. At d3, the time around which maximal destabilization was expected, the entire cell population split into three populations with respect to CD11b: Sca1^HIGH^/CD11b^LOW^ (termed α), Sca1^HIGH^/CD11b^HIGH^ (β) and unexpectedly, Sca1^LOW^/CD11b^VERY-LOW^ (γ) (Fig 3A). Single-cell transcript analysis suggests that the α-subpopulation corresponds to the destabilized but not yet fully committed cells because it displays highest cell-cell diversity and high correlation of the gene vectors (Fig 3B). The cells of subpopulation β were most advanced toward the myeloid lineage (high expression of Gfi1, CEBPα and cJun transcripts) consistent with the high CD11b expression, whereas cells of subpopulation γ correspond to “rebellious” cells that moved in the opposite direction and displayed erythroid gene expression patterns, including a large number of EpoR positive cells, despite treatment with GM-CSF/IL-3 (S5A-D Fig). At d6 the γ population disappeared (Fig 3A), consistent with the rebellious cells in the PCA analysis of Fig 1B. However, addition of EPO to the culture medium rescued the γ cells (Fig 3C), and to a lesser extent, the α but not the myeloid committed β cells.

**Fig 3.**
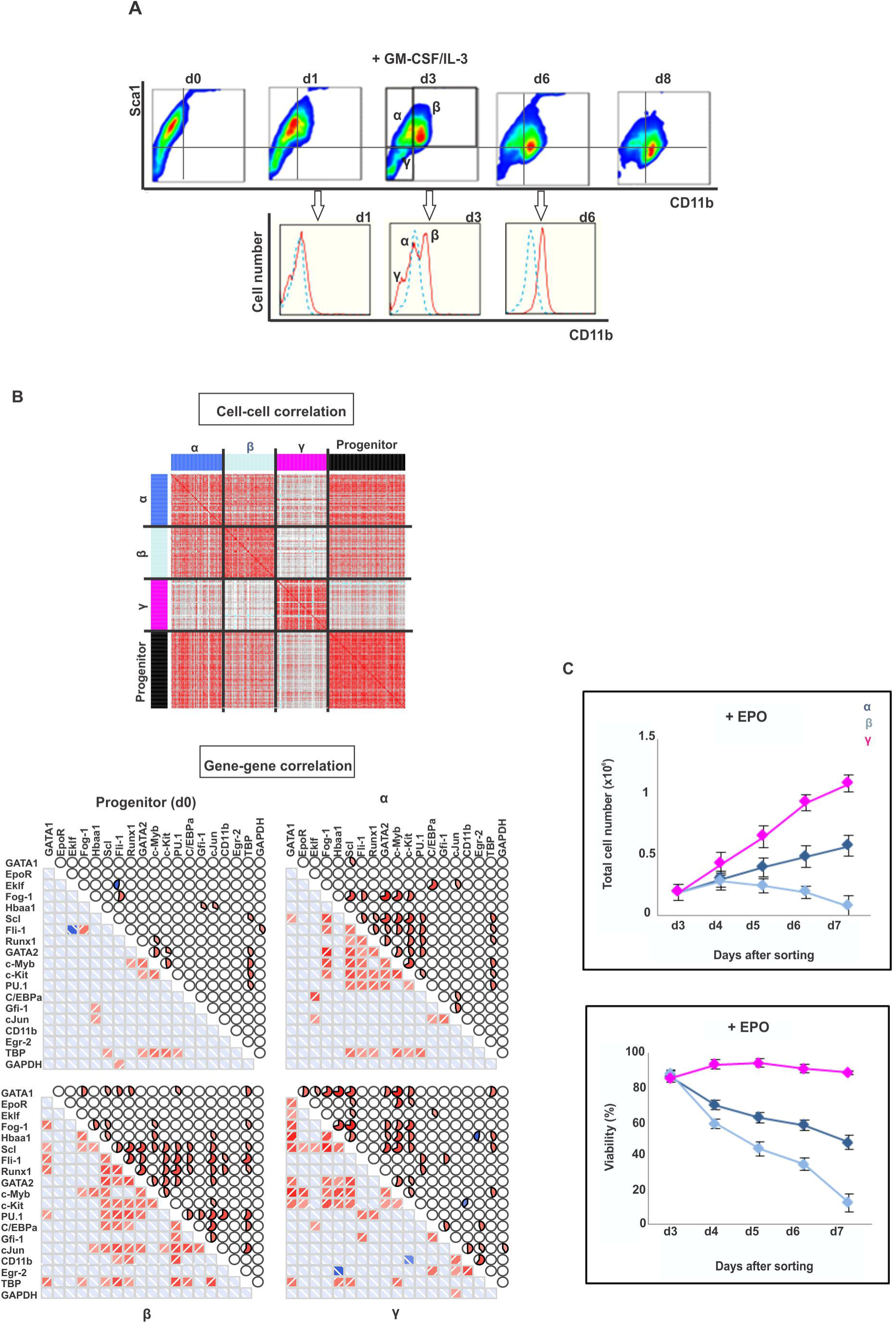
Intermediate stage of myeloid commitment exhibits destabilization of progenitor state, alternative states and “slowing down” of relaxation. **(A)** flow cytometry dot plot of expression of Sca1 and CD11b upon treatment of the progenitor EML cells with GM-CSF/IL-3. Three distinct subpopulations on d3, designated, α, β and γ, in the (tri-modal distribution of CD11b flow cytometry histogram underneath (red line, treated; blue line, untreated). **(B)** Cell-cell correlation for 72 progenitor cells and 48 cells from each of the α, β and γ subpopulations, and gene-gene correlation for all 17 genes of interest and two endogenous control genes. Pearson correlation coefficient displayed as heatmap, same color scheme as in Fig 2. **(C)** Rescue by EPO of the “rebellious” =unintended γ subpopulation (pink curve) during myeloid differentiation. Three subpopulations (α, dark blue; β, light blue and γ, pink) were FACS sorted, antibodies removed and stimulated with EPO. Total cell number and viability were quantified on day of sorting (d3) and 4 subsequent days. Viability was determined based on % of cells excluding trypan blue. Each point represents average +/- STD for 2 biological replicates.

This finding not only confirms that the rebellious γ cells have aberrantly moved towards the erythroid lineage despite myeloid instruction but also corroborates the notion of “cell selection” in fate control in which growth factors determine lineage also by acting as survival and mitogenic factors for early committed cells that express the cognate receptor – in this case the EpoR [19,23,24].

### 4. Critical slowing down

A dynamical signature of an approach to a critical transition that is often used in low dimensional systems is the “slowing down” of the relaxation of a state variable back to the original attractor state due to a reduced attracting force [15,17,18] after a small perturbation or noise-driven excursion. Although critical slowing down is linked to the flattening of the attractor and inherently associated with the increase in autocorrelation of the fluctuation of the state variables, and thus, not actually an independent criterion, its experimental assessment is distinct and often practical. Here critical slowing down was exposed by measuring the relaxation of sorted “outlier” cells which were (transiently) in an extreme state with respect to the projection into just one dimension, that of Sca1. We isolated the Sca1^LOW^ tail of populations that were either treated for 1 day with GM-CSF/IL-3 to destabilize the progenitor state, or in untreated populations. As previously shown, the Sca1^LOW^ fraction re-establishes the parental distribution within 5-6 days [4]. By contrast, cells exposed to GM-CSF/IL-3 for just one day which does not yet cause significant broadening of the distribution, required at least 9 days to reconstitute the parental Sca1 expression distribution from the same tail fraction (Fig 4).

**Fig 4.**
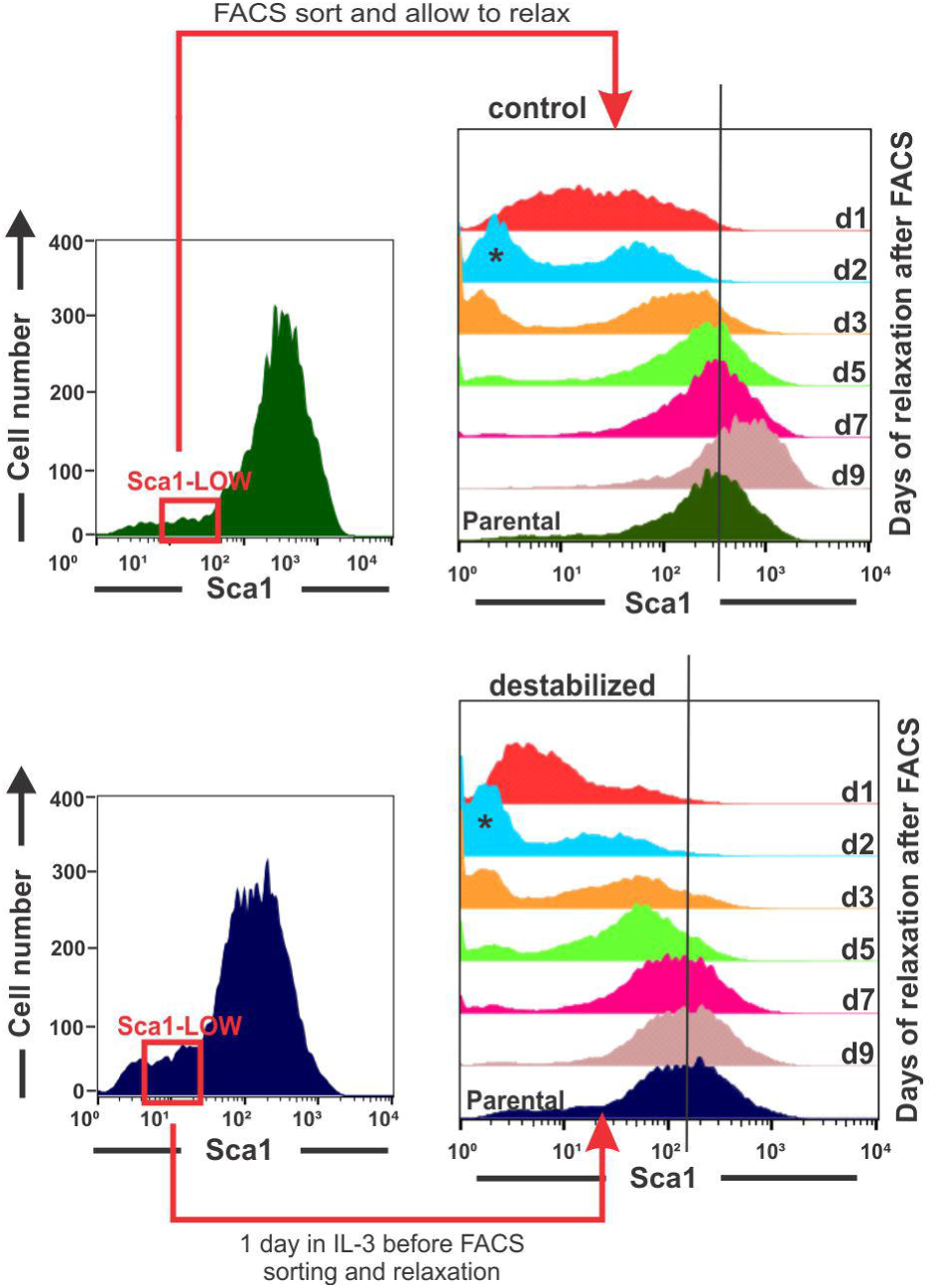
Critical Slowing down of state relaxation during fate commitment. “Critical slowing down” of relaxation and restoring of parental distribution of the sorted Sca1-low outlier fraction in the treated population. Clonal EML progenitor cells were stimulated (top) with GM-CSF/IL-3 or not (bottom) and cells with lowest 15% Sca1 expression were FACS-sorted one day after stimulation.

### 5. Transcriptomes confirm the scheme of rebellious cells

Finally, the repeated observation of “rebellious cells” is consistent with a bifurcation at which two (or more) new attractors become accessible when the progenitor attractor vanishes, representing the dichotomy between the two “sister” lineages [13,33]. The destabilization of the progenitor state, unlike in canonical saddle-node bifurcations of most studied critical transitions [15,16,17,18], opens up a choice of two attractors, and despite an instructive bias towards either one imposed by the growth factors, this allows cells to “spill” into the “wrong” attractor if molecular noise overcomes the instructive bias toward the intended lineage. Thus, the existence of “rebellious cells” is also a signature of a critical transition.

To show that such binary behavior is not an artifact of projection in one state space dimension (in this case, with respect to Sca1 or CD11b) but holds in the high-dimensional state space, we measured the transcriptomes of the subpopulations that have either responded to the growth factor or appeared to have not responded – at least with respect to change in Sca1 expression (Fig 5). As shown earlier (Fig 1) all three treatments with the either cytokines as well as combined, triggered a split of the population into two distinct subpopulations with response to the progenitor marker Sca1 (bimodal distribution at d3, Fig 5).

**Fig 5.**
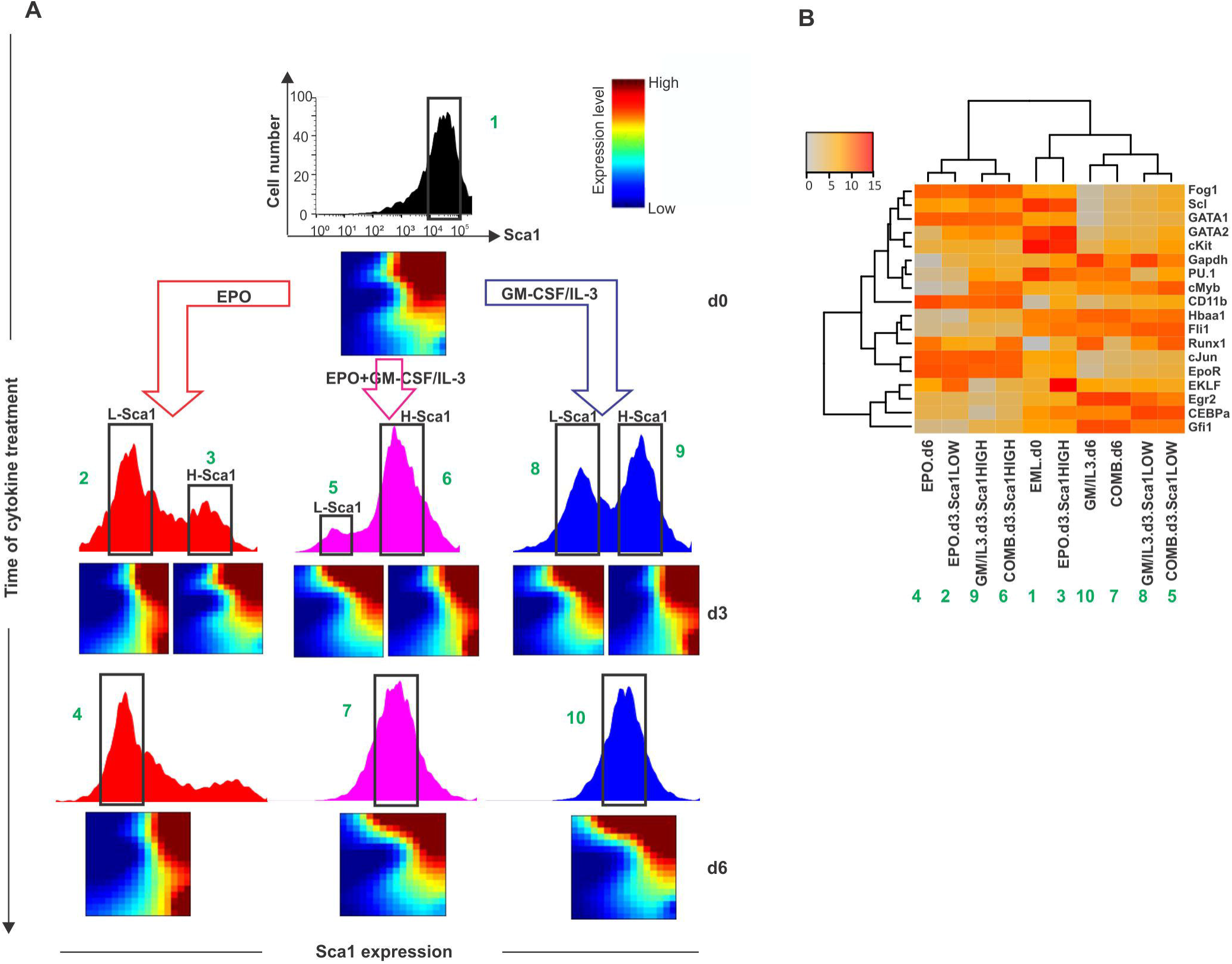
Whole-population transcriptome analysis reveals transient alternative program (“rebellious cells”). **(A)** Sca1 expression population distribution in progenitor and cytokine-treated cells and transcriptomes of sorted subpopulations at indicated treatments/time points displayed as GEDI Self-organizing maps [40]. Progenitor EML cells were stimulated with EPO alone, with GM-CSF/IL-3 alone or with the combination of the two, and the Sca1-medium (M) fractions (d0 and d6) and/or the Sca1-Low and-High subpopulations (d3) were FACS sorted and used for microarray analysis. **(B)** Hierarchical cluster analysis of the microarray-based transcriptomes of the samples in A (columns, correspondence indicated by the green numbers) for a subset of the 17 genes analyzed in single-cell qPCR (rows).

Intriguingly, cells from the Sca1^HIGH^ subpopulation which appeared to have not responded after 3d in EPO because Sca1 stayed high (fraction #3 or H-Sca1 in Fig 5A) had a transcriptome that resembled that of the cells which had responded to GM-CSF/IL-3 treatment and had down-regulated Sca1 (fraction #8 or L-Sca1 in Fig 5A). Conversely, Sca1^HIGH^ cells that had apparently not responded yet at d3 to GM-CSF/IL-3 (fraction #9 in Fig 5A) displayed a more pronounced change of the transcriptome that was remarkably similar to that of Sca1^LOW^ cells (fraction #2 that had responded to EPO). (For quantitative analysis of transcriptome similarities see S2 Table). In the combined treatment cells exhibited a transcriptome behavior that was similar to that of the nominally myeloid fated (GM-CSF/IL-3 treated) cells – in agreement with the single-cell transcript analysis (Fig 1).

The transcriptome measurement of subpopulations which appear to have not responded to the differentiation signal with respect to down-regulating the progenitor state marker actually have responded but by changes in the non-observed state space dimensions, underscoring the importance of high-dimensional dynamics. The crosswise overall similarity of the transcriptome changes in the non-responders in one treatment to that of the responders in the other treatment strongly supports the model of a constrained dynamics with a finite number (here: two) of fate options that represent the predestined developmental potentials embodied by attractors that become accessible once the progenitor state is destabilized. This behavior of aberrant but defined behavior also reveals a stochastic, non-instructive component in fate determination.

Specifically, we suspect that the rebellious cells are cells that following the flattening of the progenitor attractor initiated by the external differentiation signal erroneously enter the “non-intended” attractor that is also accessible because the stochastic gene expression fluctuations may, in some cells, overcome the instructive signal that bias the change toward a specific lineage attractor. Nevertheless the rebellious cells, being in the “wrong” fate, should eventually die because the lack of survival signals provided by the continuing presence of the same growth factor, as their disappearance in the measurement in Fig. 1 implies. Thus instruction and selection synergize, in a two-step scheme, in that cells must be instructed and be selected for in order to adopt a particular phenotype. This two-step process increases fidelity of fate determination in the tissue.

### 6. Conclusion

Here we show that exit from the multipotent progenitor state and commitment to a particular cell lineage exhibit signatures of a critical state transition because of the underlying destabilization of a high-dimensional attractor state. Fig 6 summarizes schematically the model. In doing so we confirm that the two classical models of cell fate control, instruction by extrinsic signals and selection of intrinsically predestined states [19,20,21,22,23], not only coexist but also complement each other within a formal concept.

**Fig 6.**
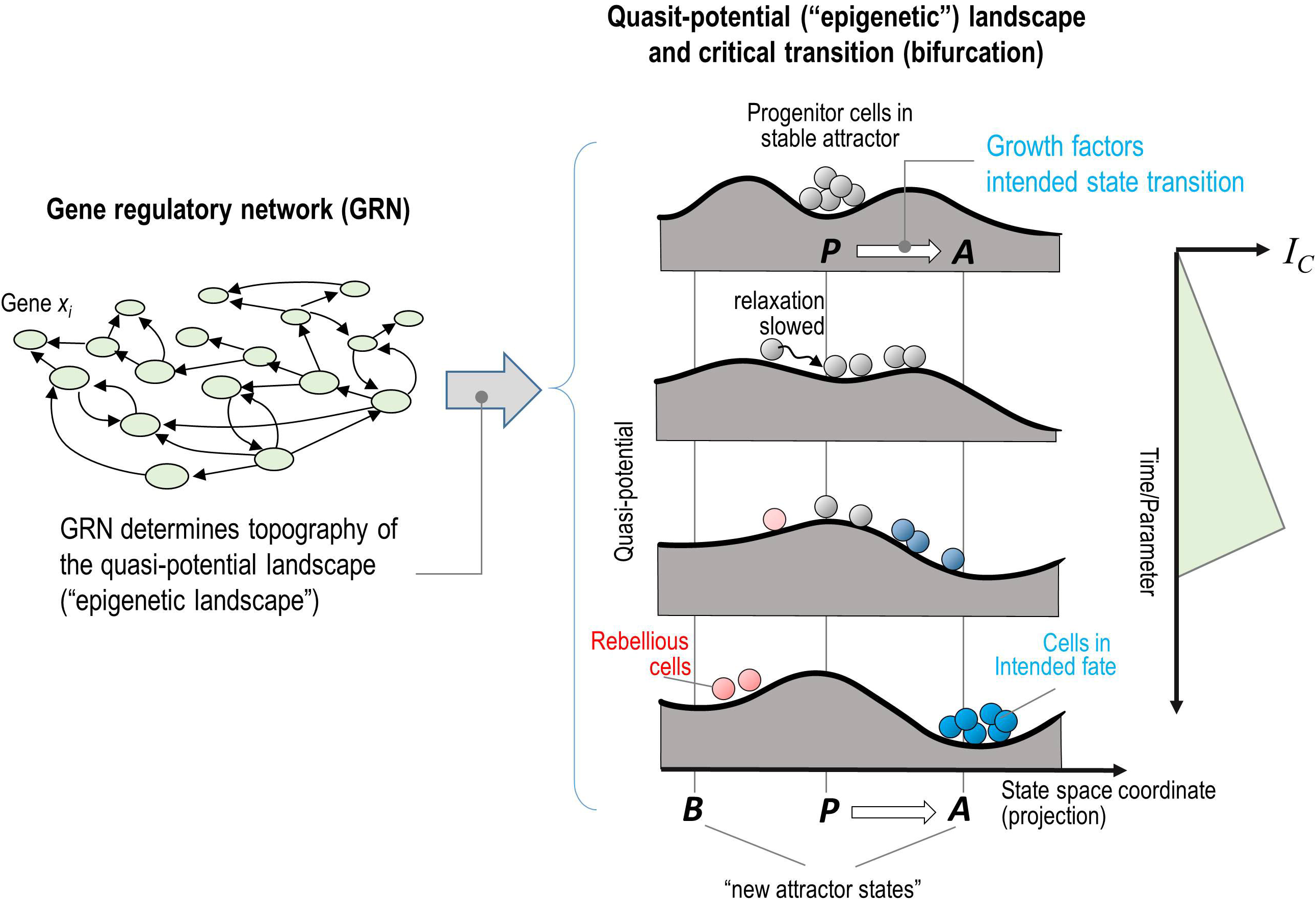
Epigenetic landscape model of symmetry-breaking bifurcation event. Progenitor cells (grey-ish) stimulated with growth factors (e.g. ATRA/IL-3). This scheme illustrates the two stages of the model: starting with the treatment of progenitor attractor state, first, the destabilization of the (meta)stable attractor of the progenitor cells and generation of a poised unstable state and second, the opening of the access to the destination attractors (both intended and non-intended), allowing the cells to descend – further instructed by the cytokines to favor one of the two valleys. As explained in section 2 of results, the cell-cell and gene-gene correlation give rise to a gradual increase of the index IC that peaked at critical transition and coincides with lineage separation in state space.

The framework of a critical transition has been used to describe sudden qualitative changes in a variety of complex systems in nature [15,16,17,18] and entails the “early warning signals” that herald the transition. We show here that early warning signs which essentially manifest the distortion of the attractor landscape that is intrinsically linked to most types of bifurcations (“tipping point”) can also be defined and detected for high-dimensional dynamics.

To do so we introduce an index *I*_*C*_, which is formally derived from dynamical systems theory [30] and whose increase serves as an early warning signal, indicating an approach to a bifurcation. *I*_*C*_ is particularly useful for single-cell resolution snapshots of molecular profiles, as provided by RNA-seq [34] and CyTOF [35], of statistical ensembles of cells (=cell populations) taken at multiple time intervals during a biological time course. This quantity is derived from a dynamical systems theory treatment of the actual underlying process and not from descriptive statistical pattern recognition as currently used to analyze single-cell molecular profiles. *I*_*C*_ captures the information immanent in both the *m* gene vectors (the expression level of a gene across a large number *n* of individual cells) and the *n* cell vectors (the state of a given cell with respect to a large number of *m* genes), resulting in the data structure of a *n* x *m* matrix for each time point in the process being studied. Thus, *I*_*C*_ does not require continuous monitoring as in many studies of critical state transitions because much of the information is in the high dimensionality (*m*) and in the statistical ensemble (*n*) and thus could be of practical utility for predicting major shifts in cell populations and tissues relevant in development and disease.

## Material and Methods

### Culture and differentiation of EML cells

Blood progenitor EML cells (ATCC CRL-11691) were cultured and maintained as described previously [24]. Multipotent EML cell population was stimulated with either EPO (to differentiate into erythroid cells), or GM-CSF/IL-3 and ATRA (to obtain myeloid cells) or a mixture of all cytokines for the “combined” treatment as previously reported [4,36]. Wright-Giemsa staining was performed with some modification following a reported protocol [37]. In brief, 60,000 cells in 250 μl of PBS + 1% FBS buffer were cytospun at 350 rpm for 5 minutes per slide and allowed to air dry for 10 minutes. Slides were subjected to five 1-second dips in methanol, followed by Wright-Giemsa staining solution (0.4% (w/v), Sigma). After a final rinse with water, slides were allowed to air dry for 30 minutes. Colored phase contrast images were obtained using a Zeiss Axiovert 200M microscope.

### Flow cytometry and fluorescent-activated cell sorting (FACS)

Cell surface protein immunostaining and flow cytometry measurements were performed using established methods [4]. Briefly the antibodies Sca1-PE (BD Pharmingen #553335), ckit-FITC (BD Pharmingen #553355) and CD11b-FITC (BD Pharmingen #557396) were used at 1:1,000 dilutions in ice-cold PBS containing 1% fetal calf serum with (flow cytometry) or without (FACS) 0.01% NaN3. Appropriate unstained and single-color controls were used for gate definition and compensation set-up. Isotype control antibodies (BD Pharmingen #553988 for FITC and #553930 for PE isotype) were used to establish the background signal caused by non-specific antibody binding. Propidium iodide (Roche #11348639001) staining was used to identify dead cells that were removed from analyses. Flow cytometry analysis was performed on a BD FACSCalibur cell cytometer with 10,000 viable events for each sample. Data were acquired using CellQuest Pro (BD) software and analyzed in FlowJo.

For FACS sorting, the Sca1 protein distribution was measured and the expression distribution was gated into three regions according to the Sca1 expression level as Sca1-Low, Mid and High on day 0, 1 and 6 or 4 regions on day 3 after differentiation initiation (Fig 1A). Single cell sorting was conducted on a BD Biosciences FACSAria III in lysis buffer (see below). For myeloid differentiation, cells were stained with antibodies for both Sca1 and CD11b protein markers and cell subpopulation were gated as illustrated in Fig 3A. For studies involving the dynamics of sorted subpopulations, antibodies were removed after sorting using brief incubation in a low-pH buffer [4].

### Single-cell gene-expression analysis using OpenArray qPCR

Single cells were directly sorted into 5.0 μl of lysis buffer (CellsDirect kit, Invitrogen) containing 4.25 μl Resuspension Buffer and 0.25 μl Lysis Enhancer using a FACSAria III (BD Biosciences). 0.5 μl RNaseOut (Invitrogen) was added to the lysis solution to protect the RNA from degradation. To ensure that liquid droplets containing single cells were deposited at the center of the well and not at the wall, the position was checked on the plastic film covering the PCR plate. To reduce the possibility of cell sticking to the wall of the PCR well plate, we used low-binding PCR plates (Axygen, #6509). As control sample, a small population of 100 cells were sorted into a single well for qPCR analysis. To test for contamination of sorted cells with mRNA from lysed dead cells, 5.5 μl liquid from the FACS instrument was collected and analyzed. After sorting, the samples were heated 75 °C for 10 min to accelerate the lysis process and samples were stored at −80 °C. From these single-cell lysate samples, cDNA was directly synthesized as described previously [36]. The obtained cDNA was pre-amplified by 18 cycles [36] and subsequently diluted with Tris-EDTA buffer at a ratio 1:10 resulting in templates for the real-time PCR analysis. This protocol led to less than 30 quantification cycles (C_q_) during the single-cell qPCR analysis on an OpenArray system (Life Technologies). On this system, each qPCR plate consists of 12×4 subarrays and each subarray contains 8×8 reaction chambers of 33 nl volume [38] (S 6A Fig). Each sample was divided into a subarray with 64 reaction chambers prior qPCR quantification. No-template (water) control was also run on each plate to check for non-specific products and/or presence of contaminants in the master mix. Following the amplification, the corresponding curves and C_q_ values were processed using the OpenArray Real-Time qPCR Analysis software (version 1.0.4) with a quantification threshold of 100(+/-5). Specific PCR primers were pre-immobilized in the chambers (S 6B Fig) and released in the first cycle by heat. For each subarray, 2 μl of target sample was loaded into each well of a 384-well plate (Applied Biosystems); subsequently, 3 μl of the master mix reaction consisting of TaqMan OpenArray Real-time PCR Master Mix (Applied Biosystems) was added to each well. Target and master mix were combined, centrifuged, and the 384-well plate was processed in the OpenArray AccuFill system (Applied Biosystems). During processing, 2.1 μl of the reaction solution was transferred automatically from each well into the corresponding subarrays of a qPCR plate, where the reaction solution retains into the reaction wells due to the differential hydrophilic–hydrophobic coating between wells and surface of the qPCR array [38]. The qPCR step was performed using thermocycling conditions of 50 °C for 2 min, 95 °C for 10 min, 40 cycles of 95 °C for 15 sec and 60 °C for 1 min.

### Testing Taman qPCR assays

We used off-the-shelf primers designed by Applied BioSystems (Life Technologies) for the analysis. The primers are usually designed to span exon-exon junction to target multiple splice variants of one transcript and to target only and specifically the gene of interest, avoiding amplification of genomic DNA. S3 Table lists all genes of interest, the inventoried TaqMan assay IDs (Applied Biosystems) and further relevant information where the manufacturer does not provide primer and probe sequences. To evaluate qPCR assay performance, calibration (standard) curves were generated by performing qPCR on a serial dilution of a prepared template. Each of these dilutions was dispensed into two subarrays of OpenArray plate leading to 6 technical qPCR replicates for each single cell sample. To minimize the effect of sampling errors on quantification precision, only sample/assay combinations with at least 3 quantifiable replicates were considered for preparing the standard curves. The GAPDH assay was not pre-immobilized on OpenArray plate but was independently tested on BioRad qPCR platform.

### Analysis of single-cell gene expression data

Data analysis is described in more details in Supplementary Discussion. Single-cell expression data were initially analysed with OpenArray qPCR analysis software. For quality control, amplification curves were quality filtered and Ct thresholds were set for each assay with the same thresholds used across all experiments and cell populations. Data were subsequently exported to Excel as csv files. All of Cq values are available in S1 Table. Samples not expressing any gene were excluded from the analysis. Experimentally determined LODs were used as cutoff Cqs (S3 Table). Each assay was performed in triplicates, and the median of the triplicates was used for subsequent analysis. After this pre-processing, ΔCq was calculated as previously described [39]. Higher level of analysis such as correlation, clustering, and PCA was performed on log2-transfromed expression data.

### Gene expression profiling with microarrays and data analysis

Microarray analyses were performed by the Vancouver Prostate Cancer centre. EML progenitor cell population was stimulated with EPO alone, IL-3/GM-CSF alone or a combination of all cytokines. On d3 and d6 after stimulation with different cytokines, the main “peaks” in the Sca1 distribution were gated and cell subpopulations were sorted using FACSAria III. Fig 1A and B illustrate the experimental design for the microarray experiments. Total RNA was extracted from 1×10^6^ of sorted subpopulations using mirVana miRNA Isolation Kit (Ambion) following the manufacturer’s instructions. Genomic DNA was removed from the isolated and purified RNA using DNase I. Total RNA quality was assessed with the Agilent 2100 Bioanalyzer prior to microarray analysis. Samples with a RIN value equal to or greater than 8.0 were deemed acceptable for microarray analysis. Samples were prepared following Agilent’s One-Color Microarray-Based Gene Expression Analysis Low Input Quick Amp Labeling v6.0. An input of 100 ng of total RNA was used to generate Cyanine-3 labeled cRNA. Samples were hybridized on Agilent SurePrint G3 Mouse GE 8x60K Microarray (Design ID 028005). Arrays were scanned with the Agilent DNA Microarray Scanner at a 3 μm scan resolution, and data was processed with Agilent Feature Extraction 11.0.1.1. To filter out genes that were not expressed above the background noise, a raw intensity cutoff value of 25 was applied because the correlation between the technical replicates decreases for higher levels. Green processed signal was quantile-normalized using the “normalize.quantiles” function in R that takes care of inter-chip variability. To filter out genes which did not change between the samples, the distribution of each gene across all samples was analyzed. Therefore the standard deviation (STD) distribution was calculated and only genes with STD > 10% were selected. As a result, 6297 genes passed the criteria and were selected as the 10% top genes among the samples. Self-organising maps (SOM) of the 10% top most varied genes (6297 genes) were generated using the Gene Expression Dynamics Inspector program (GEDI) [40]. Cluster analysis was performed using the “clustergram” function in Matlab R2012a Bioinformatics toolbox using hierarchical clustering with Euclidean distance metric and average linkage to generate the dendrogram. Input data was log2-tranformed values of normalized fluorescent intensity signals for genes of interest extracted from the samples and plotted as a heatmap. Data represented the average of *n* = 2 independent biological replicates. The normalized fluorescent intensity values of 17 genes of interest in the curated network were extracted from each sample.

## Acknowledgments

The authors would like to thank Drs. Luonan Chen and Hong Qian for helpful discussions.

## Author contributions

M.M. designed experiments, performed experiments and data analysis. A.S. designed and performed statistical data analysis and theoretical analysis. J.Z. and A.G. performed theoretical analysis, R.L-Q and I.G.C. and H.C. performed experiments. S.H. conceived of experiments and theory and designed experiments and conceived and performed theoretical analysis. S.H. drafted the manuscript, M.M., A.S. and S.H. edited and wrote the paper.

## Additional information

Accession codes: Microarray data have been deposited in GEO under accession number GSE70405.

## Supporting Information Captions

**S1 Appendix. Supplementary methods (data analysis)**

**S2 Appendix. Supplementary discussion (with mathematical proof)**

**S1 Fig. Manually curated model of gene regulatory network governing fate decision of CMP**. Network of experimentally verified regulatory interactions of transcription factors involved in multipotency of the CMP state, fate decision and differentiation to the erythroid and myeloid lineages (S1 Table). The canonical GATA1-PU.1 circuit is highlighted in green. A few surface markers including c-kit (progenitor, grey box), EpoR (erythroid, red box) and CD11b (myeloid, blue box) were included in the network to control the cell differentiation behavior and used as markers for lineage commitment in experiments. The numbers point to the references listed in S1 Table.

**S2 Fig. Gene expression profile of single-cell samples during differentiation**.

Expression profiles of 17 transcription factors and control genes (rows) in individual cells (columns) are visualized as a heatmap. Cell columns are arranged for days d1, d3 and d6 with respect to different treatments where grey shades correspond to untreated progenitors (d0), red shades to EPO treatment, blue shades indicate cells treated with GM-CSF/IL-3 and purple shades to combined treatment EPO+GM-CSF/IL-3 cytokines. The different shades of each color indicate the different Sca1 marker expression levels Sca1^Low^ (L), Sca1^Mid^ (M) and Sca1^High^ (H) determined during FACS sorting where darker shades denote higher Sca1 expression. Gene rows were ordered according to their biological role as indicated on the left.

**S3 Fig. Technical noise associated with single-cell RT-qPCR is significantly smaller than biological cell-cell variability. (A)** Quantification cycles (Cq) of 80 individual EML cells for GATA1 expression is reported. Values are means ± STD for up to 128 technical replicates. **(B)** Quantification cycles (Cq) of up to 110 technical replicates are presented for 3 selected single-cells. Single-cell Cqs of biological samples clearly show a broader distribution relative to that of technical replicates. **(C)** Box plots represent the variability in terms of CV for technical replicates averaged over 110 realizations of the real-time PCR-steps on the ds-cDNA and the distribution of CV across all 80 individual EML progenitor cells for the GATA1 expression. The biological variation was significantly larger than the technical noise (p-value 2.2e-28, Mann-Whitney U test). Similar results were obtained for PU.1 (not shown).

**S4 Fig. Distinct trajectories of cell differentiation are observed upon stimulation of progenitor cells with cytokines in the PCA state space. (A)** Principal component projections in a total of ~1600 haematopoietic cells including progenitor (black), single-EPO treated (red-shades), single-IL3/GM-CSF treated (blue-shades) and combined-treated (purple-shades) in the first three components determined from expression of all 17 transcription factors and endogenous control genes. **(B)** Principal component loadings for PC 2 and 3 indicate the extent to which each gene contributes to the separation of cells along each component. **(C)** PCA weights of genes for the first three PCs reveals the importance of the individual genes to explain the difference between the different treatments and corresponding cell fate. **(D)** Cells in their attractor states still exhibit heterogeneous transcription profiles that can be traced back to individual genes. Cells treated with GM-CSF/IL-3 for 6 days are clearly located within the state space defined by the myeloid genes and cells treated by EPO exhibit 2 clusters where the lower one is governed by erythroid genes and the higher one by stemness genes. **(E)** Variance explained by principal components show that the first three components jointly explain more than 70% of variation in the data.

**S5 Fig. Gene expression in individual cells from the progenitor population and the α, β, and γ subpopulations. (A-D)** Heatmap representation of gene expression profiles for the set of 17 genes of the curated network and 2 endogenous genes as control in total 216 single cells including 72 progenitor cells (panel A) and 48 single cells from each of the three subpopulations in the tri-modal Sca-1 population distribution on day 3 after GM-CSF/IL-3 treatment (Fig 3), α (B) β (C), and γ (D). Genes are ordered according to their reported biological role, as erythroid-associated (red box), stemness (green box), myloid-associated (blue box) and endogenous genes in all subplots. Based on the expressed genes, the β subpopulation seems to be committed to the myeloid lineage while the γ subpopulation is committed to the erythroid lineage. The α subpopulation exhints an indeterminacy with a bias towards the myeloid lineage. **(E)** PCA of all attractor cells (d0 and d6) as shown in the S4 Fig combined with the cells from the α (yellow), β (green), and γ (pink) subpopulations support the above described similarity to the untreated EML, the GM-CSF/IL-3 stimulated and the EPO-stimulated cells, respectively. **(F)** Coefficient of variation CV (used as a cell-specific quantity to expose population dispersion) was calculated for each cell from the expression levels across all genes for each subpopulation. Histograms represent the number of cells at different level of the CV measure and show that cells in α subpopulation have higher spread of cellular CV values.

**S6 Fig. Representation of an OpenArray plate used for single-cell qPCR. (A)** Each OpenArray (Applied Biosystems) is the size of a microscope slide. It holds 48 groups (subarrays, red rectangular) of 64 holes of 33 nl volume in which one PCR reaction occurs. A hydrophilic layer is at the interior surface of each hole and a hydrophobic layer is at the exterior surface of the plate allowing for filling the hole by surface tension. In total, each array carries 3072 qPCR reactions. **(B)** Specific PCR primers are pre-immobilized in individual holes (by manufacturer, for customized assay patterns) and released by heat in the first cycle. **(C)** An example of the distribution of single-cell samples (SC) along with NTC (no template water control), IRC (inter-run calibrator) and 100-cell control (PC) samples on an OpenArray chip.

**S7 Fig. Quality control of single-cell qCPR. (A)** Inter-chip variability is evaluated using inter-run calibrator (IRC) sample. Each curve represents the distribution of Cq values of each gene across all OpenArray chips. The flat black curve represents the distribution of all genes across all chips. The inter-gene differences are up to 2 orders of magnitude larger than the inter-chip variability of the same gene. The inter-run calibrator was a 10-fold diluted sample of 18 cycles pre-amplified cDNA of 10 ng isolated RNA from EML progenitor cell population. **(B-D)** Correlation between gene expression in an ensemble of 48 individual cells and 6 replicates of 100-cell pools is plotted. Cells used were from subpopulations, α, β and γ (subplots b-d) as presented in Fig 3 and 19 genes as listed in Table S3 were measured in triplicate in all single cells and bulk (100-cell) samples from each subpopulation. Mean expression for each gene was calculated across all single cell or pool samples. Note that the scaled mean expression for 100-cells pool was plotted against mean expression for single-cells. In all cases a high correlation between single-cell data and bulk data with correlation coefficient of > 0.86 was observed.

**S1 Table. Regulatory interactions in the curated GRN model of binary fate decision in CMP**. Table of the regulatory interactions (either activating (A) or inhibiting (I)) between the genes. For each interaction, the literature is referenced. All interactions have been reported in for murine hematopoiesis.

**S2 Table. Quantified dissimilarity between transcriptomes from micro-arrays** between samples. Pair-wise dissimilarity between expression profiles (samples) was calculated based on the normalized gene expression levels for 6297 filtered genes (see METHODS) with 1 – *R* where *R* is the Pearson’s correlation coefficient which ranges from 0 to 1, meaning that 0 correspond to highest similarity and 1 to most different expression. Bootstrapping was performed by randomly selecting 30% of the genes in any sample to calculate the pair-wise dissimilarity metric and repeating the procedure 10,000 times to generate the reported standard deviations.

**S3 Table. Evaluation of qPCR assays**. Table lists all primer pairs and relevant information including IDs and amplicon length. All assays were inventoried. Identical PCR primers were used in the pre-amplification step and the subsequent singleplex qPCR step. In addition, the amplification efficiency and limit of detection (LOD) of the qPCR assays are given. To evaluate efficiency and LOD, a 1:2 serial dilution was prepared from 18 cycles pre-amplified product from 10 ng RNA purified from EML progenitor cell population. Amplification efficiency was calculated according to: [10(1/-S)-1] × 100%. The slope was obtained by linear regression of the standards curve. Efficiency was determined as average of two biological replicates with 6 qPCR technical replicates each. The Cq value for the LOD is defined as the most diluted sample that results in positive amplification for 5 out of 6 replicates.

**S4 Table: Single-cell and 100-cell samples quantification cycles (raw) data**. The quantification cycles (Cqs) for all analyzed single-cells as well as 100-cell-pool control samples are reported. Single cells from untreated EML control cells as well as EML cells treated with EPO, GM-CSF/IL-3 or a combination of all cytokines on d1, d3 and d6 of stimulation. Gene expression data for single-cell samples sorted from α, β and γ subpopulations generated upon GM-CSF/IL-3 treatment of EML are also included. 6 replicates of the 100-cell samples were also sorted from each fraction and/or subpopulation and analyzed as control.

## References

1. Macarthur BD, Ma’ayan A, Lemischka IR. Systems biology of stem cell fate and cellular reprogramming. Nat. Rev. Mol. Cell Biol. 2009;10:672–81.

2. Huang S. Cell lineage determination in state space: a systems view brings flexibility to dogmatic canonical rules. PLoS Biol. 2010;8:e1000380.

3. Wray J, Kalkan T, Smith AG. The ground state of pluripotency. Biochem. Soc. Trans. 2010;38:1027–1032.

4. Chang HH, Hemberg M, Barahona M, Ingber DE, Huang S. Transcriptome-wide noise controls lineage choice in mammalian progenitor cells. Nature. 2008;453:544–7.

5. Hough SR, Laslett AL, Grimmond SB, Kolle G, Pera MF, A continuum of cell states spans pluripotency and lineage commitment in human embryonic stem cells. PLoS One. 2009;4(11):e7708.

6. Pina C, Fugazza C, Tipping AJ, Brown J, Soneji S, Teles J, et al., Inferring rules of lineage commitment in haematopoiesis. Nat. Cell Biol. 2010;14:287–94.

7. Canham MA, Sharov AA, Ko MSH, Brickman JM. Functional heterogeneity of embryonic stem cells revealed through translational amplification of an early endodermal transcript. PLoS Biol. 2010;8:e1000379.

8. Kauffman S. Homeostasis and differentiation in random genetic control networks. Nature. 1969;224:177–8.

9. Wu M, Su RQ, Li X, Ellis T, Lai YC, Wang X. Engineering of regulated stochastic cell fate determination. Proc Natl Acad Sci U S A. 2013; 110:10610–5.

10. Mendoza L, Alvarez-Buylla ER. Dynamics of the genetic regulatory network for Arabidopsis thaliana flower morphogenesis. J Theor Biol. 1998;193:307–19.

11. Huang S, Eichler G, Bar-Yam Y, Ingber DE. Cell fates as high-dimensional attractor states of a complex gene regulatory network. Phys Rev Lett. 2005;94:128701.

12. Zhou JX, Aliyu MDS, Aurell E, Huang S. Quasi-potential landscape in complex multi-stable systems. J. R. Soc. Interface. 2015;12:1–15.

13. Zhou JX, Huang S. Understanding gene circuits at cell-fate branch points for rational cell reprogramming. Trends Genet. 2011;27:55–62.

14. Huang S. Systems biology of stem cells: three useful perspectives to help overcome the paradigm of linear pathways. Philos Trans R Soc Lond B Biol Sci. 2011;366:2247–59.

15. Sole RV. Phase Transitions. Princeton, NJ.;Princeton University Press; 2011.

16. Scheffer M, Carpenter SR, Lenton TM, Bascompte J, Brock W, Dakos V, et al., Anticipating critical transitions. Science. 2012;338:344–8.

17. Trefois C, Antony PM, Goncalves J, Skupin A, Balling R. Critical transitions in chronic disease□: transferring concepts from ecology to systems medicine. Curr. Opin. Biotechnol. 2015;34:48–55.

18. Scheffer M, Bascompte J, Brock WA, Brovkin V, Carpenter SR, Dakos V, et al. Early-warning signals for critical transitions. Nature. 2009;461:53–9.

19. Coffman RL, Reiner SL. Instruction, selection, or tampering with the odds? Science. 1999 284:1283–1285.

20. Germain RN. T-cell development and the CD4-CD8 lineage decision. Nat Rev Immunol. 2002;2:309–22.

21. Robb L. Cytokine receptors and hematopoietic differentiation. Oncogene. 2007;26:6715–23.

22. Graf T. Differentiation plasticity of hematopoietic cells. Blood. 2002;99:3089–101.

23. Enver T, Jacobsen SEW. Instructions writ in blood. Nature. 2009;461:183–184.

24. Enver T, Heyworth CM, Dexter TM. Do stem cells play dice? Blood. 1998;92:358–41; discussion 352.

25. Davis CB, Killeen N, Crooks ME, Raulet D, Littman DR. Evidence for a stochastic mechanism in the differentiation of mature subsets of T lymphocytes. Cell. 1993;73:237–47.

26. Metcalf D. Lineage commitment and maturation in hematopoietic cells: the case for extrinsic regulation. Blood. 1998;92:345–7; discussion 352.

27. Rieger MA, Hoppe PS, Smejkal BM, Eitelhuber AC, Schroeder T. Hematopoietic cytokines can instruct lineage choice. Science. 2009;325:217–8.

28. Waddington CH. Principles of Embryology. Allen Unwin Ltd; New York:Macmillan; 1956.

29. Wang J, Xu L, Wang E, Huang S. The potential landscape of genetic circuits imposes the arrow of time in stem cell differentiation. Biophys. J. 2010;99:29–39.

30. Chen L, Liu R, Liu ZP, Li M, Aihara K. Detecting early-warning signals for sudden deterioration of complex diseases by dynamical network biomarkers. Sci. Rep. 2012;2:18–20.

31. Giuliani A. Statistical Mechanics of Gene Expression Networks□: Increasing Connectivity as a Response to Stressful Condition. Adv. Syst. Biol. 2014;3:1–4.

32. Gorban AN, Smirnova EV, Tyukina T. Correlations, risk and crisis: From physiology to finance. Phys. A Stat. Mech. its Appl. 2010;389:3193–3217.

33. Huang S, Guo YP, May G, Enver T. Bifurcation dynamics in lineage-commitment in bipotent progenitor cells. Dev. Biol. 2007;305: 695–713.

34. Bandura DR, Baranov VL, Ornatsky OI, Antonov A, Kinach R, Lou X, Pavlov S, Vorobiev S, Dick JE, Tanner SD. Mass Cytometry: technique for real time single cell multitarget immunoassay based on inductively coupled plasma time-of-flight mass spectrometry. Anal. Chem. 2009;81:6813–6822.

35. Ozsolak F, Milos PM. RNA sequencing: advances, challenges and opportunities. Nature Reviews:Genetics. 2011;12:87–98.

36. Mojtahedi M, Fouquier d’Hérouël A, Huang S. Direct elicitation of template concentration from quantification cycle (Cq) distributions in digital PCR. Nucleic Acids Res. 2014;42:e126.

37. Woronzoff-Dashkoff KK. The wright-giemsa stain. Secrets revealed. Clin. Lab. Med. 2002;22:15–23.

38. Branon C, Morrison T. Nanoliter high throughput quantitative PCR. Nucleic Acids Res. 2006;34:e123.

39. Fluidigm. Application Guidance□: Single-Cell Data Analysis-RevA1. 2012; 1–40.

40. Guo Y, Eichler GS, Feng Y, Ingber DE, Huang S. Towards a holistic, yet gene-centered analysis of gene expression profiles: A case study of human lung cancers. J. Biomed. Biotechnol. 2006;29:1–11.

